# Stimulus content shapes cortical response statistics

**DOI:** 10.1101/238774

**Authors:** Mihály Bányai, Andreea Lazar, Liane Klein, Johanna Klon-Lipok, Wolf Singer, Gergő Orbán

## Abstract

Spike count correlations (SCCs) are ubiquitous in sensory cortices, are characterized by rich structure and arise from structured internal interactions. Yet, most theories of visual perception focus exclusively on the mean responses of individual neurons. Here, we argue that feedback interactions in primary visual cortex (V1) establish the context in which individual neurons process complex stimuli and that changes in visual context give rise to stimulus-dependent SCCs. Measuring V1 population responses to natural scenes in behaving macaques, we show that the fine structure of SCCs is stimulus-specific and variations in response correlations across-stimuli are independent of variations in response means. Moreover, we demonstrate that stimulus-specificity of SCCs in V1 can be directly manipulated by controlling the high-order structure of synthetic stimuli. We propose that stimulus-specificity of SCCs is a natural consequence of hierarchical inference where inferences on the presence of high-level image features modulate inferences on the presence of low-level features.

## Introduction

Spike-count correlations (SCCs), covariation of neuronal responses across multiple presentations of the same stimulus, are ubiquitous in sensory cortices and span different modalities (Downer et al., 2015; Petersen et al., 2001; Romo et al., 2003) and processing stages (Cohen and Maunsell, 2009; Nienborg and Cumming, 2006; Ponce-Alvarez et al., 2013; Zohary et al., 1994). In the visual system, SCCs, also termed noise correlations, have traditionally been considered to be independent of the stimulus and hence to impede stimulus encoding (Averbeck et al., 2006). Studies on stimulus-independent aspects of SCCs in the primary visual cortex (V1) sought to capture correlation patterns that were solely accounted for by differences in receptive field structure (Ecker et al., 2010; Gutnisky and Dragoi, 2008). Initial investigations of stimulus-dependence of SCCs focussed on the mean of SCCs (Cohen and Kohn, 2011; Kohn and M. A. Smith, 2005) but stimulus-dependent changes in the mean are modest in awake animals (Ecker et al., 2010; Rikhye and Sur, 2015). However, recent studies using calcium imaging of V1 in awake mice revealed a dependence of the fine structure of correlations on stimulus-statistics (Hofer et al., 2011; Rikhye and Sur, 2015), suggesting that not only mean responses (first order statistics) but response correlations (second order statistics) too, could carry useful stimulus-specific information.

SCCs reflect the internal dynamics of the network and specific feedback interactions were shown to contribute to their emergence in a number of ways (Ecker et al., 2016; Lin et al., 2015; Rabinowitz et al., 2015; Rosenbaum et al., 2016). In V1, feedback interactions comprise both lateral connections from the local circuitry and top-down connections from higher level areas (Harris and Mrsic-Flogel, 2013). Investigations into the organization of feedback revealed that characteristic regularities present in the visual environment are reflected in the structure of both lateral (Kaschube, 2014; Löwel and Singer, 1992; McGuire et al., 1991; Schmidt et al., 1997; G. B. Smith et al., 2015) and top-down (Klink et al., 2017; Lee and Nguyen, 2001) interactions. These feedback interactions could thus establish dependencies between the responses of neurons whose receptive fields are sensitive to stimulus features that have high probability to co-occur in the visual environment and support perceptual grouping, such as the vicinity, continuity, collinearity, color and direction of motion of composing scene elements. Structured feedback interactions are expected to introduce correlated variability in neuronal responses. Indeed, patterns in lateral connections that reflect receptive field similarity of neurons (Cossell et al., 2015) were shown to capture the stimulus-independent patterns in SCC structure (Rosenbaum et al., 2016). Importantly, this intricate feedback circuitry seems to distort the sensory information carried by bottom-up pathways, which is also reflected in the recurrence of activity patterns of spontaneous activity during activity evoked by visual stimulation (Arieli et al., 1996; Berkes et al., 2011; Fiser et al., 2004). The observation that the structure of correlations reflects the structure of feedback and that the structure of feedback reflects the statistics of environmental stimuli raises the intriguing prospect that correlations and their modulations can provide important insights into the computations performed by coordinated populations during visual perception.

Natural visual stimuli are complex and provide insufficient information for unambiguous interpretation. Evidence suggests that the visual system represents an internal model of the environment, which serves the integration of information about the current stimulus with previously acquired knowledge of natural scene statistics (Berkes et al., 2011; Fiser et al., 2010; Frégnac and Bathellier, 2015; Weiss et al., 2002; Yuille and Kersten, 2006). In the process of perceptual inference, the internal model contributes information about the expected activation and coactivation structure of neurons (Orbán et al., 2016) and provides a context for the interpretation of sensory input (Coen-Cagli et al., 2015; Yuille and Kersten, 2006). Crucially, when interpreting complex stimuli, the context plays an essential role: the likelihood of the presence or absence of a particular visual feature is dependent on the presence or absence of a large number of contextual features. Indeed, on a pebbly beach we expect different co-occurrence patterns of elementary edges, characteristic to V1 simple cell receptive fields, than on a wheat field. Furthermore, given that the visual cortex processes visual information in a series of hierarchical processing stages, contextual information from the higher levels of the processing hierarchy can inform and constrain the activity at lower levels of processing through feedback (Gilbert and Sigman, 2007; Klink et al., 2017; Lee and Mumford, 2003). For instance, merging information from an extended area in the visual field will inform low-level stages of the processing hierarchy that we are on a beach and the expected textures are characterized by specific spatial frequencies and lines are characterized by specific curvatures. Thus, we predict that in the primary visual cortex, where neurons are sensitive to simple features like oriented edges, high-level visual context constrains the internal dynamics though specific feedback, and ultimately results in stimulus-specific SCC structure.

Feedback modulation of higher order statistics of responses, including variance (Goris et al., 2014) and correlations (Ecker et al., 2014) in V1 were shown to contribute to multiplicative effects in activity fluctuations. Indeed, patterns in V1 SCCs in response to periodic grating stimuli were shown to be aligned with a simple phenomenological model of V1 responses which only considers stimulus dependence of neuronal responses in terms of tuning curves and assumes that joint modulations are stimulus-independent (Lin et al., 2015). This model, however, does not aim to account for stimulus-dependent modulations in feedback and therefore does not predict stimulus-specificity of SCCs. Similarly, a functional account of V1 which links (co)variability of neuronal responses to perceptual uncertainty but lacks a representation of higher-order stimulus features fails to predict stimulus-specificity in the correlation structure (Orbán et al., 2016). Modulations of the fine structure of correlations have been predicted by functional models of attention (Haefner et al., 2016), which related changes in correlation structure to changes in task variables. Indeed, attentional modulations have long been considered as a major source of top-down influences (Gilbert and Sigman, 2007; Reynolds and Heeger, 2009), which affect not only the mean of neuronal responses, but the correlations as well (Cohen and Maunsell, 2009; Rabinowitz et al., 2015; Ruff and Cohen, 2014). Here we go beyond these accounts and argue that hierarchical perceptual inference has a direct predictable effect on the structure of spike count correlations in V1, that is independent from task or attentional influences. We hypothesize that, for stimuli with high-order structure, inferences on the presence of high-level visual features modulate inferences on the presence of low-level features through specific feedback to V1 neurons, leading to stimulus-specificity in SCC patterns. Conversely, without high-level structure stimulus-specificity of correlation patterns is expected to dwindle.

To test this hypothesis, we designed an experiment in which we can characterize the full correlation matrix, the so called partial correlations (Yatsenko et al., 2015). First, we established that correlation patterns in response to natural images are stimulus-specific. We developed the contrastive rate matching method to identify modulations in correlation structure that are independent of changes in the mean of the responses. Next, we designed synthetic image families with low-level or high-level structure. These image families are distinguished by the level of processing hierarchy where samples from the family elicit selective responses from individual neurons. Importantly, in a hierarchical model of visual perception, high-level synthetic images, but not low-level synthetic images, are expected to elicit stimulus-specific feedback structure and consequently stimulus-specific correlations. We confirm these predictions and demonstrate that the stimulus specificity of SCCs is dependent on stimulus-structure: synthetic stimuli characterized solely by low-level structure elicit correlation patterns with reduced stimulus-specificity, while synthetic stimuli characterized by high-level structure restore stimulus-specificity of correlations.

## Results

In order to phrase predictions on the effect of stimulus-specific feedback interactions on SCCs, we introduce a hierarchical model of visual processing in the ventral stream. The model naturally extends earlier probabilistic models of V1 activity (Coen-Cagli et al., 2012; Olshausen and Field, 1996; Orbán et al., 2016; Schwartz and Simoncelli, 2001) by assuming an additional layer of processing. The additional layer is analogous to higher processing layers in the ventral stream and, for simplicity, we identify it with the secondary visual cortex (V2). V2 neurons are assumed to be selective to texture like patterns (Freeman et al., 2013) that emerge from combinations of elementary features (i.e. Gabor filters). Probabilistic models of perceptual inference, similar to the one proposed here, have been motivated by the fundamentally noisy and ambiguous nature of environmental stimuli, which gained extensive experimental support from behavioral studies (Kersten et al., 2004; Schwartz et al., 2009; Weiss et al., 2002). Importantly, probabilistic computations suggest that uncertainties about the inferred environmental features need to be maintained by an efficient system and therefore we consider neural representations which can represent such uncertainties (Hoyer and Hyvarinen, 2003; Lee and Mumford, 2003; Orbán et al., 2016).

Assuming a hierarchical internal model for the representation of natural images in the visual cortex (Fig 1A), probabilistic inference in the model corresponds to stimulus perception (Helmholtz, 1962). In this context, activities of neurons correspond to activation of variables, and selectivities of neurons correspond to filter properties of variables. In this model, the activity level of a neuron is assumed to represent the inferred intensity of its preferred visual feature and at different levels of the hierarchy, neurons are sensitive to features of different complexity. In a widely used approximation, the receptive fields of V1 neurons can be characterized by Gabor filters, while the receptive fields of V2 neurons can be characterized by texture-like filters (Freeman et al., 2013). When stimuli are noisy or ambiguous, the model incorporates knowledge about the uncertainty associated with inference at different levels of the visual hierarchy in the form of a posterior distribution. Upon the presentation of a particular image, x, the posterior distribution for the activations of V1 neurons, y, conveys detailed information about the uncertainty of the features represented by V1 receptive fields, including the specification of not only mean activations but variances and covariances as well (see also Supplementary Note for detailed derivation):

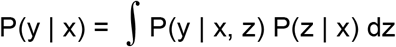

**Figure 1.**
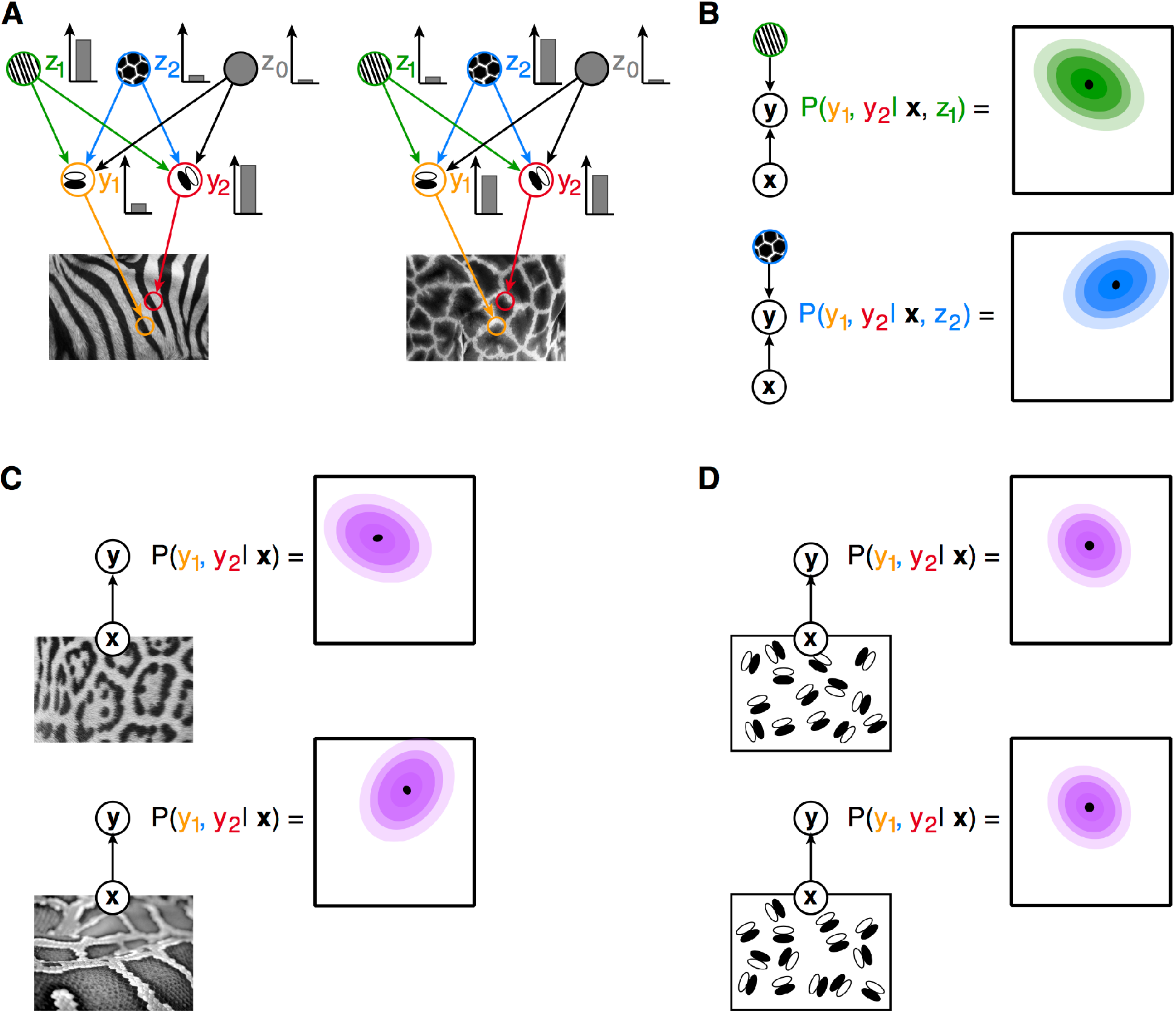
Illustration of inference in a hierarchical statistical model. (**A**) An image, **x**, is assumed to be generated by combining features of different complexity: high-level features, z_i_ (green and blue circles), determine the large-scale structure of low-level features, e.g. textures determine the joint statistics of edges (z_0_ is a bias term that represents an interpretation where no higher-order structure is present). Low-level features, y_i_, capture simple regularities in images, e.g. darker and lighter image areas underlying edges (orange and red circles). In the visual system, upon presentation of a stimulus, the contribution of different features to the observed image is inferred: different images (left and right panels) elicit different intensity responses from the neurons (inset bar plots). (**B**) The statistical internal model establishes a joint probability distribution for the coactivation of low-level features upon observing a stimulus: beyond the most probable joint activations (black dots) a wide range of co-activations is compatible with the high-level percept, albeit with different probabilities (colors matching those on panel **A**). Given the activation of a particular high-level feature (z_1_ or z_2_ for the left and right panels, respectively), thejoint distribution over activations of low-level features (contours) displays a covariance specific to the high-level feature. (**C**) The posterior distribution for low-level features is characterized both by the mean and covariance of the distribution (top panel). Posterior distribution for a distinct structured image (bottom panel) is characterized by a different mean and correlation structure. (**D**) A stimulus with no higher level structure is invoking an interpretation that low-level features are independent, therefore the correlation structure of images with only low-level structure will be identical. Arrows define conditional dependencies throughout the figure.

The first term of the integral is the probability distribution of the joint activations of V1 neurons given a particular image and a particular set of activations, z, at a hierarchical level beyond V1 (Fig. 1B). The second term establishes weights for averaging over possible high-level activations. This equation highlights three important points: 1, Activations at the lower level of the hierarchy, V1, depend on high-level activations, i.e. specific predictions can be obtained for top-down interactions; 2, Activations at V1 can be correlated, i.e. if a high-level feature represented in V2 assigns high probability to particular combinations of features then variability in z will induce correlations in y (Figs. 1B); 3, Since the probability of different combinations of high-level activations, P(z | x), changes with changes in the stimulus, correlations in V1 will be stimulus-dependent. As a consequence, hierarchical statistical inference predicts stimulus-dependent correlations for structured stimuli, e.g. for natural images, thus reflecting top-down influences (Fig. 1C). However, in the absence of high-level structure stimuli will not be informative with respect to high-level inferences and therefore will result in unspecific top-down influences and hence, unspecific correlations (Fig. 1D). Anatomical connections that contribute to the implementation of probabilistic computations in the hierarchical internal model are expected to involve not only bottom-up and top-down projections but lateral connections as well. These connections are essential for implementing the local interaction patterns between V1 neurons, thus contributing to the nonlinear interaction patterns of receptive fields (Kaschube, 2014; Schmidt et al., 1997; G. B. Smith et al., 2015).

### Stimulus-dependence of spike count correlations

Parallel multielectrode recordings (32 channels) were obtained from area V1 of two awake behaving monkeys (macaca mulatta). The receptive fields of the recorded units were located approximately at 3° (Monkey A) and 5° (Monkey I) from the fixation spot. Monkeys were trained to perform an attention task in which, after initiating fixation (Fig. 2A), they were presented with a pair of natural images at two locations, left and right from the fixation spot, one of which overlapped with the RFs of the recorded units. After 700 ms, a change in fixation spot color cued the monkeys to report an incoming change in either the left or right image. The task was used to ensure the engagement of the animal and our analysis was constrained to neural responses evoked by stimulus presentation before appearance of the cue signal (see details of the task in the Experimental Procedures). The initial transient responses after stimulus onset were omitted from the analysis to reduce stimulus locked correlations, leaving a window of 400 ms to assess response statistics (Fig. 2A). Reliable estimation of the full spike count correlation matrix between recorded channels required a large number of repetitions, therefore the number of different images was limited to 6 or 8 images per session, providing a range of 65-180 repetitions per image. Mean response, as characterized by the firing rate, was selective for stimulus identity (Fig. 2A), which is captured by a high level of dissimilarity of firing rate patterns in response to a range of different natural images compared to lower dissimilarity of responses to identical stimulus presentations (Fig. 2B).

**Figure 2.**
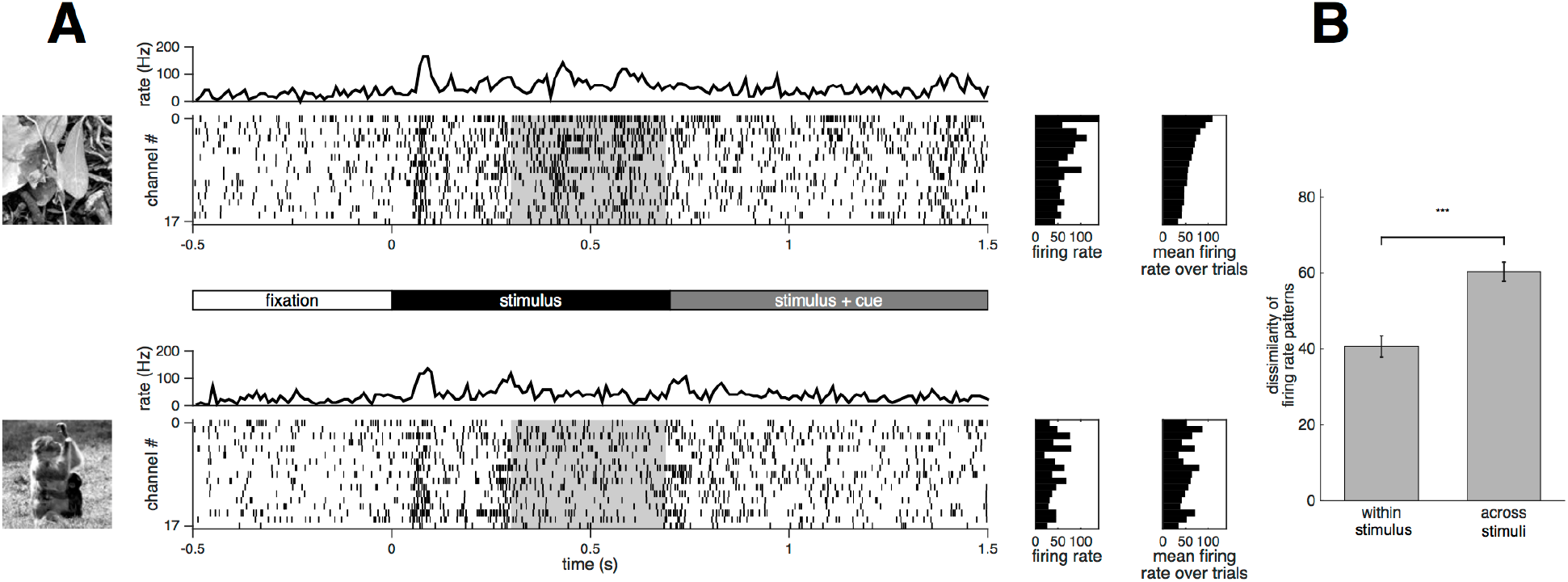
Structure of the experiment and mean neural responses. **a**, Time course of neural responses upon the presentation of two natural images (top and bottom panels). After presenting a fixation point for 500 ms (timeline in the middle), a pair of stimuli are presented off-foveally at equal distances from the fixation point. One of the images (shown on the left for the example trial) covers the receptive fields of recorded V1 neurons. After another 700 ms the color of the fixation point changes, cuing the monkey which of the images it needs to focus its attention to. In the following 800 ms one of the images is rotated, and the monkey is asked to respond if the cued stimulus changes and to withhold responses to changes of the non – cued stimulus. Multiunit activity is recorded on multiple channels and spiking activity is obtained (raster plots). Stimulus onset elicits large transient responses (peaks in the channel-averaged activity, top trace), followed by more sustained but weaker responses. Analysis of spiking activity was constrained to this segment of 400 ms (gray shading). Mean activity levels differ on different channels in a single trial (left side raster plot) but reflect the overall pattern of activation levels measured across trials where the same image was presented to the neurons (right side raster plot). A different image (bottom panel) elicits different mean activations, as reflected by the individual trial and by trial-averaged responses. Ordering of the channels is preserved across the two panels and ordering was done according to the trial-average responses to the first image (top-right raster). **b**, Dissimilarity of patterns in average firing rates calculated across half of the trials are compared to average firing rates calculated for the rest of the trials for the same stimulus (within-stimulus) or to average firing rates measured for other stimuli (across-stimuli).

First, our goal was to establish the stimulus-specificity of the fine correlation patterns in population responses to natural image patches. For each stimulus, we calculated a spike count correlation matrix, by extracting correlations between the activities of any two neurons across repeated presentations of the same stimulus (Fig. 3A). We analyzed the stimulus-specificity of the structure of SCC matrices by comparing the difference between the correlation matrices extracted in two different conditions: 1, from two independent subsets of data in response to the same stimulus (within-stimulus); 2, from the responses of neurons to different stimuli (across-stimuli, Fig. 3B and Fig. S1B). This treatment goes beyond traditional approaches that only characterize the population mean of the distribution of correlations (Fig. 3C). Measurement of SCCs from a finite number of trials is noisy and therefore estimates of the SCC matrix are variable (Fig. 3A). As a consequence, the within-stimulus difference of SCC matrices is finite and this difference can be used to establish a baseline for the estimates of stimulus-specificity of SCCs. This baseline shrinks with increasing number of trials (set size). To establish the number of trials needed for a reliable estimate of spike count correlations, we assessed dissimilarity as a function of set size (Fig. 3D). There is a steep drop in dissimilarity at low trial counts due to high variance of correlation estimates. We balanced the trade-off between the number of repetitions and the size of the stimulus set in an experimental session by aiming for approximately 80 repetitions per stimulus.

**Figure 3.**
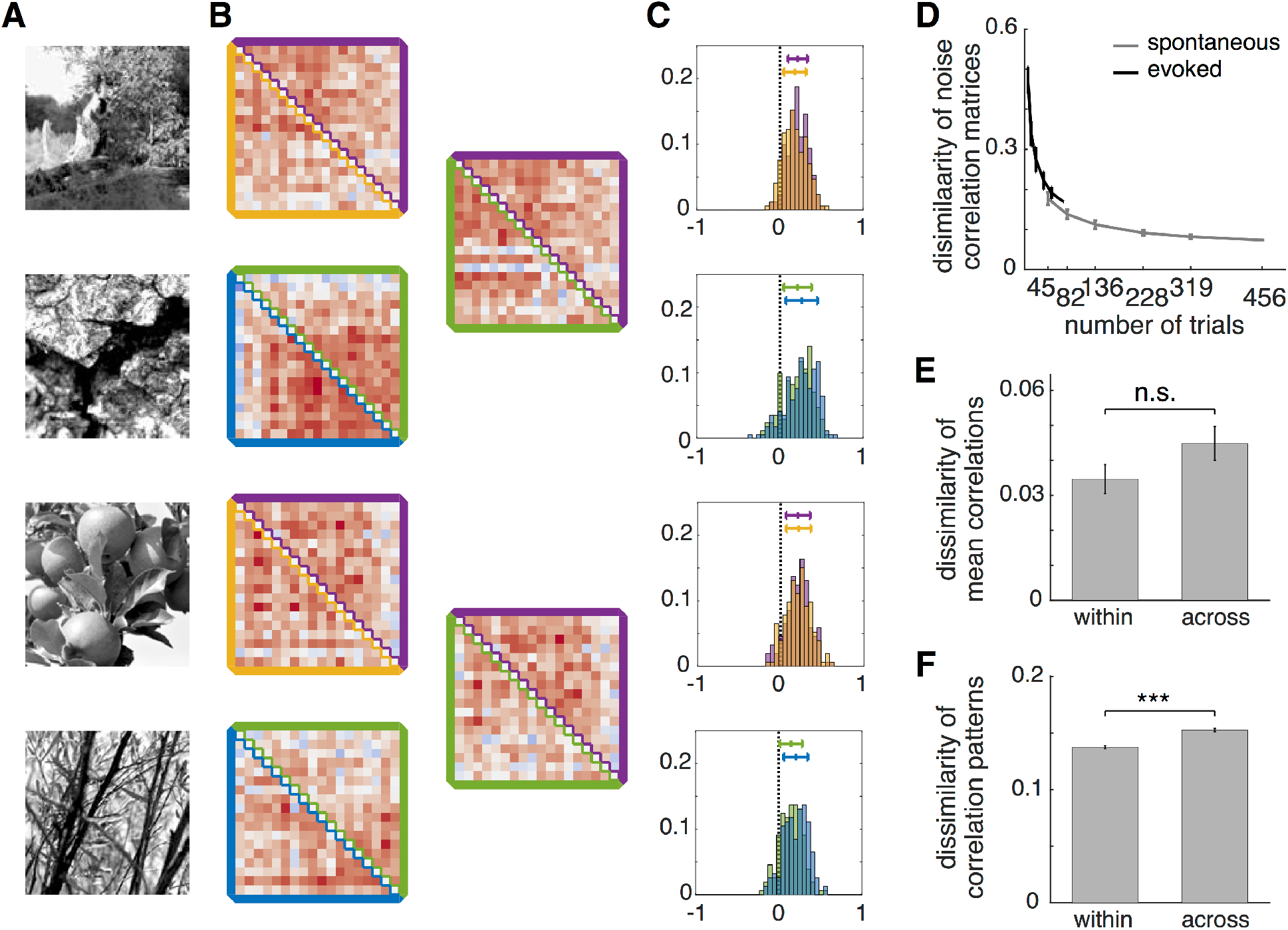
Stimulus-dependence of spike count correlation. **a**, Natural images elicit responses that are characterized by spike count correlations. **b**, Fine structure of spike count correlations (partial correlations) can be determined for the two subsets of trials (upper and lower triangles of the spike count correlation matrix, with colors matching those used on **a**), and they can be compared to both sets of trials coming from responses to the same image (left column, within-stimulus), or to different images (right column, across-stimuli). **c**, Histograms of spike count correlations calculated between channels are obtained for two equal-sized subset of trials (two different colors on the same plot). Means of the correlations are close to zero but the distribution has a large spread (tick and whisker plots with matching colors). **d**, Dependence of within-stimulus dissimilarity of spike count correlation matrices on the number of trials used to estimate pairwise spike count correlations. **e**, Dissimilarity of mean spike count correlations within-stimulus and across-stimuli. **f**, Dissimilarity of spike count correlation matrices within-stimulus and across-stimuli.

We checked whether stimulus-specificity of SCCs can be established based on the mean of the correlation distribution. Comparison of changes in the mean was not conclusive since the dissimilarity of the mean correlation across stimuli was not significantly higher than that within stimulus (t-test, p=0.12, t=-1.59, df=82, Fig. 3E). Comparison of SCC matrices instead of the mean of SCC distributions is sensitive to changes in the patterns of correlations and therefore provides a more detailed information on the stimulus-dependence of population responses (Fig. 3F). Dissimilarity of SCC matrices was significantly higher across stimuli than within stimulus (t-test, p=5.6e-14, t=-7.53, df=12258). We also determined that the significance of the difference in dissimilarities is not merely the result of a larger sample size due to the large number of elements of correlation matrices. To this end we constructed a measure that matches the sample size of the population mean of correlations. We calculated a single dissimilarity value for a particular pair of stimuli and compared this measure across conditions (t-test, p=3.7e-4, t=-3.71, df=82; Fig. S1C). Comparison of correlation matrices implicitly establishes a comparison between two multivariate normal distributions. A widely used measure to assess the dissimilarity of probability distributions is the Kullback-Leibler (KL) divergence, which can be calculated analytically for Normal distributions and can be used to assess the dissimilarity of the correlation structures. We found a similar pattern in the difference in dissimilarities with KL divergence as with other measures (t-test, p=1.52e-3, t=-3.28, df=82, Fig. S1D). Taken together, these analyses indicate that fine patterns in spike count correlations are specific to natural stimuli.

### Contrastive rate matching

Firing rate has a major effect on spike count correlations estimated from spiking activity (de La Rocha et al., 2007) and is one of the major factors that affect our analyses (Schulz et al., 2015) (see also Experimental Procedures). As a consequence, firing rate changes could constitute a confound for establishing stimulus-specificity of spike count correlations. To eliminate this potential confound we designed a method, contrastive rate matching, to control for the effect of firing rate changes on spike count correlation estimates (Experimental Procedures, Fig. 4). Briefly, in a given condition the distributions of changes in firing rate and correlation were calculated (Fig. 4A). In the two-dimensional distribution every data point represents a pair of channels: the magnitude of mean firing rate difference is plotted against the magnitude of change in spike count correlation. A similar distribution of firing rate and correlation differences was constructed for the condition with which correlation changes are contrasted (Fig. 4B). To eliminate the dependence of the estimate of correlation change on firing rate changes, the marginal distribution of firing rate changes is matched across the two conditions by subsampling the data points. On these subsampled distributions of firing rate and correlation differences, the magnitude of firing rate changes will be equal in the two conditions and the residual condition-dependence of correlations can be assessed.

**Figure 4.**
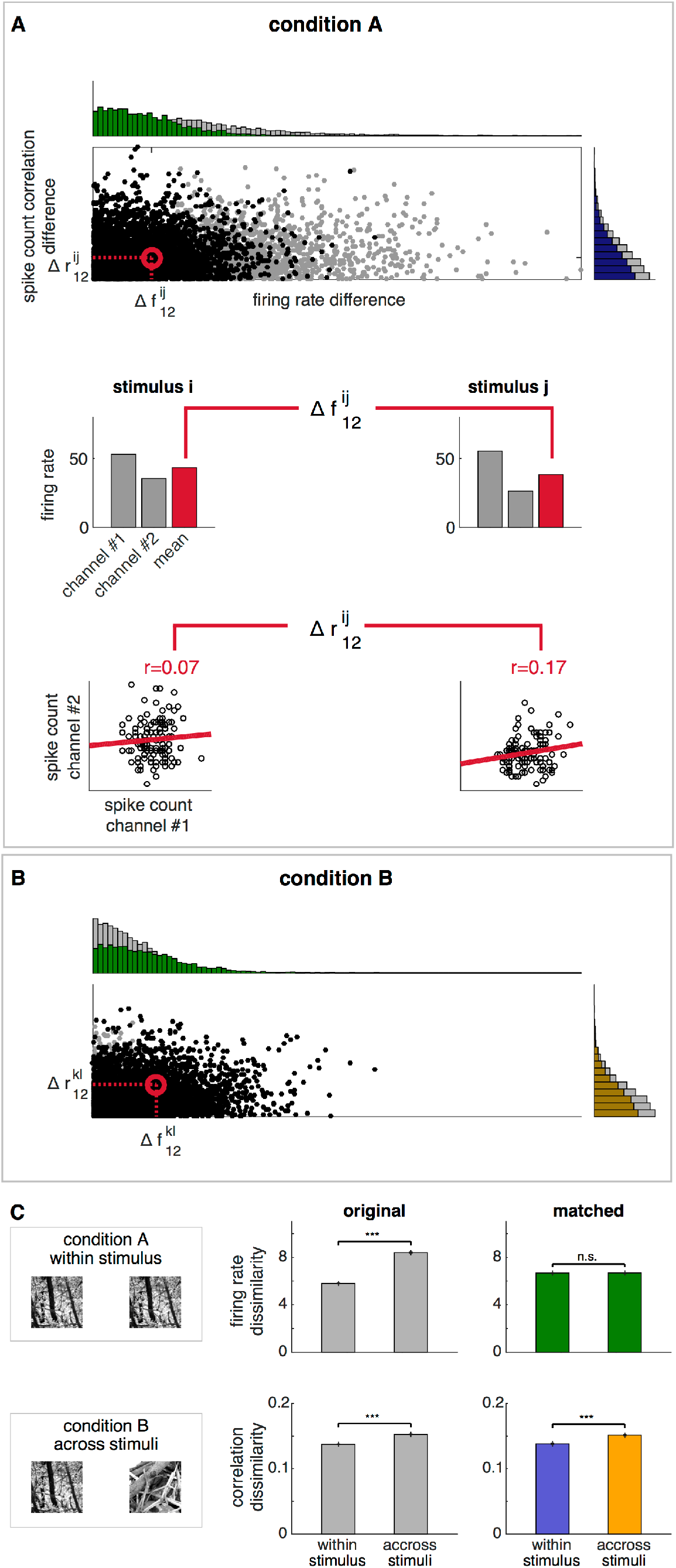
Controlling changes in firing rate distribution for the comparison of spike count correlation distributions. **a**, Distribution of the magnitude of changes in firing rates and correlations upon presenting stimulus ‘i’ or ‘j’. Every point on the scatter plot represents the response of a pair of channels to every pair of stimuli in a given condition (grey dots). Data from all sessions are aggregated. Marginal distributions for firing rate differences and correlation differences are shown on the horizontal and vertical histograms, respectively. To calculate the difference in firing rates, a mean response is calculated for the two channels at the two stimuli (grey bars on middle panels), next geometric mean is obtained (red bars on middle panels) and finally the absolute value of difference is calculated. To calculate the spike count correlation difference of the same pair of cells, z-scored spike counts are obtained (bottom panels, representing individual trials), Pearson correlation is calculated for each image and finally the difference between correlations calculated for the two images is calculated. **b**, Under a different condition where stimuli ‘k’ and ‘l’ are presented, joint distribution of firing rate differences and correlation differences are calculated (grey dots). Since we want to assess correlation changes independent of changes in firing rates, the marginal distributions of firing rate differences in condition A (a) and condition B (b) are subsampled such that firing rate differences have the same distribution under the two conditions (green histograms). Firing rate difference-matched correlation differences are obtained by calculating correlation difference distributions (gold and purple histograms) from the subsampled joint distributions (black dots on the scatter on both a and b). c, Using within-stimulus comparison as condition A and across-stimuli comparison as condition B (left panel), dissimilarity of firing rates (top row) and correlations (bottom row) when firing rate differences are not equated (middle panels) or matched (right panels). Initial differences in firing rate dissimilarity (top row, grey bars) are eliminated by the matching procedure (top row, green bars). Initial differences in the dissimilarity of correlations (bottom row, grey bars) remain significant after matching firing rate differences (bottom row, colored bars, colors matching those of histograms at panels **a** and **b**).

To demonstrate the power of contrastive rate matching we used synthetic data in which the two conditions can be fully controlled (Fig. S2). A network of 40 neurons was simulated in which membrane potential correlations and firing rates were set for each condition and the simulation matched the experimental conditions in terms of the amount of data used. In each experiment the first condition assessed had identical firing rate and spike count correlation profiles in every trial (Fig. S2A). We investigated three different scenarios for the second condition. First, dissimilarity of spike count correlation matrices was assessed across trials with identical spike count correlation patterns but different mean activations (Fig. S2B). Under these conditions we expect that due to firing rate differences, spike count correlations will vary across trials with different firing rates. Indeed, dissimilarity of correlation matrices is higher in the condition where firing rate differences are present even though the membrane potential correlations are identical (Fig. S2E). However, contrastive rate matching eliminates this difference. Second, dissimilarity of spike count correlation matrices was assessed across trials with identical mean activations but different spike count correlation patterns (Fig. S2C). As expected, dissimilarity of correlations remained significant in both the non-matched and in the matched cases (Fig. S2F). The last analysis tested the scenario where both firing rates and correlations show differences across trials (Fig. S2D). Residual differences in correlation dissimilarity after contrastive rate matching demonstrated that differences in membrane potential correlations can be identified in spike count correlations (Fig. S2G).

We assessed within-stimulus and across-stimuli dissimilarity of spike count correlation matrices using contrastive rate matching on data recorded from V1 (Fig. 4C). As expected, contrastive rate matching eliminates the condition-dependence of firing rate dissimilarity (t-test, p=0.95, t=-0.06, df=9918). Also, the analysis confirmed that differences in correlation dissimilarity are significant even after contrastive rate matching (t-test, p=3.5e-9, t=-5.91, df=9918), therefore stimulus-specificity of the fine structure of SCC is not a result of changes in firing rates.

### Stimulus-structure dependence of spike count correlations

We argued that higher-order structure in stimuli elicits differential responses at the network level both in V1 and at higher levels of processing. We identified stimulus-specific correlation structure in V1 activity as a result of differential feedback from higher levels of processing to V1 neurons. This prediction, however, is not an exclusive consequence of hierarchical inference since it can be accounted for by other models. Therefore we designed an experiment which exploits the selectivities of neurons at different levels of the processing hierarchy to control stimulus-specificity of SCCs and formulated a more specific prediction: if stimulus-specificity is a consequence of stimulus-specific feedback from higher levels of processing then removing higher-order structure from images should reduce stimulus-specificity of correlations.

We tested this hypothesis explicitly by recording additional electrophysiological data in two monkeys, performing the same task. In these experiments, we interleaved natural images with synthetic images, composed of independent Gabor filters, that retained the low-level structure prefered by individual V1 simple cells but contained no further dependencies (low level (LL)-synthetic stimuli, see also Experimental Procedures). We calculated the average dissimilarity of firing rates and spike count correlation matrices across natural images and compared them to the average dissimilarity of firing rates and correlations across LL-synthetic stimuli (Fig. 5A). We found that both firing rate dissimilarity and correlation dissimilarity was significantly higher for natural images than for LL-synthetic stimuli (Fig. 5B, t-test, p=7.4e-286, t=37.29, df=10474 and p=1.46e-21, t=9.56, df=10474 for firing rate and correlation, respectively).

**Figure 5.**
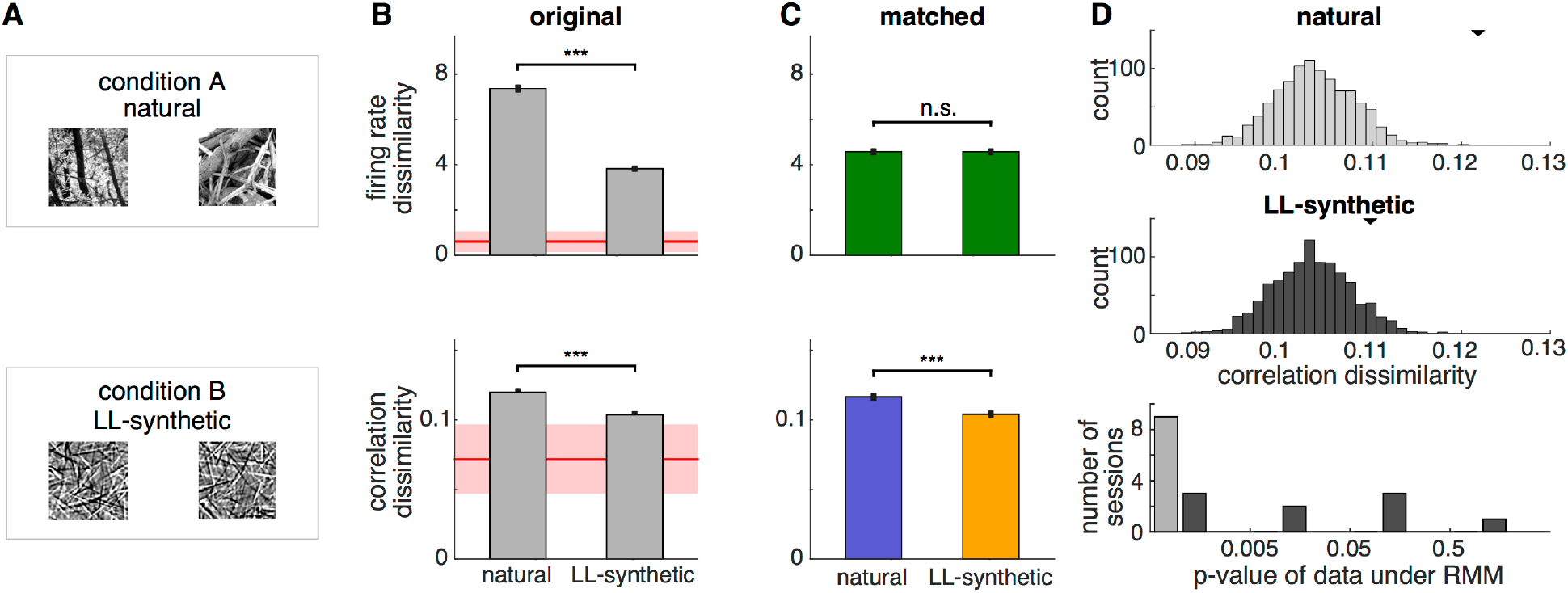
Comparison of stimulus-specificity of correlation patterns induced by different stimulus structures. **a**, A set of natural image patches are used as a reference condition and a set of synthetic image patches generated from a V1 model of images is used in the second condition. **b**, Stimulus-specificity of firing rate responses (top panel) and spike count correlation patterns (bottom panel) in the original (unmatched) data. While correlations show higher specificity for natural images, specificity of firing rate responses is also higher in the reference condition. Shaded areas show the extrapolated estimate of within-stimulus dissimilarity for both firing rates and correlations (see Experimental Procedures). **c**, Contrastive rate matching eliminates stimulus-specificity of firing rate responses, but the residual dissimilarity of spike count correlations is still significantly higher for natural images than for LL-synthetic stimuli. **d**, Raster marginal models (RMMs) fitted to the spike trains recorded under natural image stimulation condition and under LL-synthetic stimulus stimulation condition in an example session (top and middle, respectively). Distributions of dissimilarities are calculated between correlation matrices sampled from RMMs obtained from the population activities recorded for individual stimuli. Black triangles mark the mean dissimilarity calculated from the electrophysiological data. Bottom: Likelihoods of recorded dissimilarity under the RMM model in all of the sessions in the natural and LL-synthetic conditions (colors match those of the top and middle panels). Dissimilarity indices of 500 pairs of correlation matrices sampled from the RMM model were used to assess the likelihood of the recorded data. Stimulus-dependence of correlation matrices under natural image stimulation could not be explained by an RMM model in any of the recorded sessions. Dissimilarity determined for LL-synthetic stimuli was significantly different from the RMM model in five out of nine sessions.

The limited number of available repetitions establishes a lower bound on the dissimilarity measures (Fig. 5D). To directly obtain a lower bound for this experiment, data would be required to be split into two halves and within-stimulus dissimilarities should be computed across the split data. Such a manipulation, however, would result in higher variance in our primary measure of interest, the across-stimuli dissimilarity. Therefore we obtained the lower bound indirectly, by extrapolating within-stimulus dissimilarity from dissimilarities calculated for lower numbers of repetitions (by subsampling the available data, see also Experimental Procedures).

Altering stimulus content induces variations in firing rates and these variations can in turn affect correlation measures. If stimulus specificity of firing rates were higher in V1 for natural than synthetic images correlation dissimilarity would be affected by firing rate changes. Therefore we determined stimulus-statistics specificity of correlation dissimilarity using contrastive rate matching (Fig. 5C) which eliminates differences between spike count correlations caused by firing rate dissimilarity (t-test, p=0.99, t=0.015, df=7468). After this correction (removal of confound) residual spike count correlation dissimilarity was still much higher for natural than for LL-synthetic stimuli and highly significant (t-test p=4.17e-10, t=6.26, df=7468).

Since our measurements are based on a finite population, firing rate has an effect on the variability of measured correlations: higher firing rates can result in a lower number of possible binary combinations formed from spikes. To control for this possible confound we constructed surrogate data from a phenomenological model of population activity, the raster marginal model (RMM, Experimental Procedures). The RMM provided a distribution of correlation matrices for every single image. Correlation dissimilarities were calculated from 1000 correlation matrices obtained from each distribution. The histogram of correlation dissimilarities determines how likely it is that the dissimilarity measured on the data can be traced back to changes in basic firing statistics. Histograms obtained for the two conditions did not show significant differences in their mean (t-test, p=0.12, t=1.54, df=1998) but the histogram for natural images revealed that the dissimilarity obtained from the data was in the tail of the distributions of possible dissimilarities (p<0.001, while for synthetic images p=0.076; Fig. 5D). Analysis of all recorded sessions reveals that in all cases the activity evoked by natural images is highly unlikely under a simple RMM account. However, spiking responses to the LL-synthetic stimulus set were significantly different from an RMM account only in five out of nine trials at the p<0.05 level. Taken together, comparison of the results obtained with natural and LL-synthetic data, excludes the possibility that observed dissimilarities were merely resulting from changes that an RMM model can account for.

### Higher-order structure over elementary features induces stimulus-specific correlation patterns

Hierarchical inference predicts that stimulus-specificity of SCCs is a consequence of specific feedback, and it is the inference on the presence of high-level image features which modulates inference on the presence of low-level features. Neurons responsible for high-level inferences are sensitive to combinations of elementary features (Nassi and Callaway, 2009). In particular, neurons in V2 were shown to be selective to parameters that define texture-like patterns (Freeman et al., 2013). Motivated by these findings, we generated, a novel set of synthetic stimuli that combined Gabor filters into texture-like patterns, thus introducing the kind of higher-level structure, which is expected to elicit differential responses in V2.

In novel recordings, we interleaved synthetic images characterized by low-level structure (LL-synthetic stimuli) with texture-like synthetic images characterized by high-level structure (HL-synthetic stimuli). Both firing rates and spike count correlations showed higher specificity for HL-synthetic stimuli than for LL-synthetic stimuli (Fig. 6B, t-test, p=2.6e-180, t=29.3, df=8998 and p=5.56e-16, t=8.11, df=8998 for firing rates and correlations, respectively). Contrastive rate matching was applied to eliminate differences caused by firing rate dissimilarity (Fig. 6C, t-test, p=0.94, t=0.076, df=6794). This manipulation did not alter the conclusion on stimulus-specificity of SCCs: the difference between the correlation dissimilarity remained significant (t-test, p=8.55e-09, t=5.76, df=6794).

**Figure 6.**
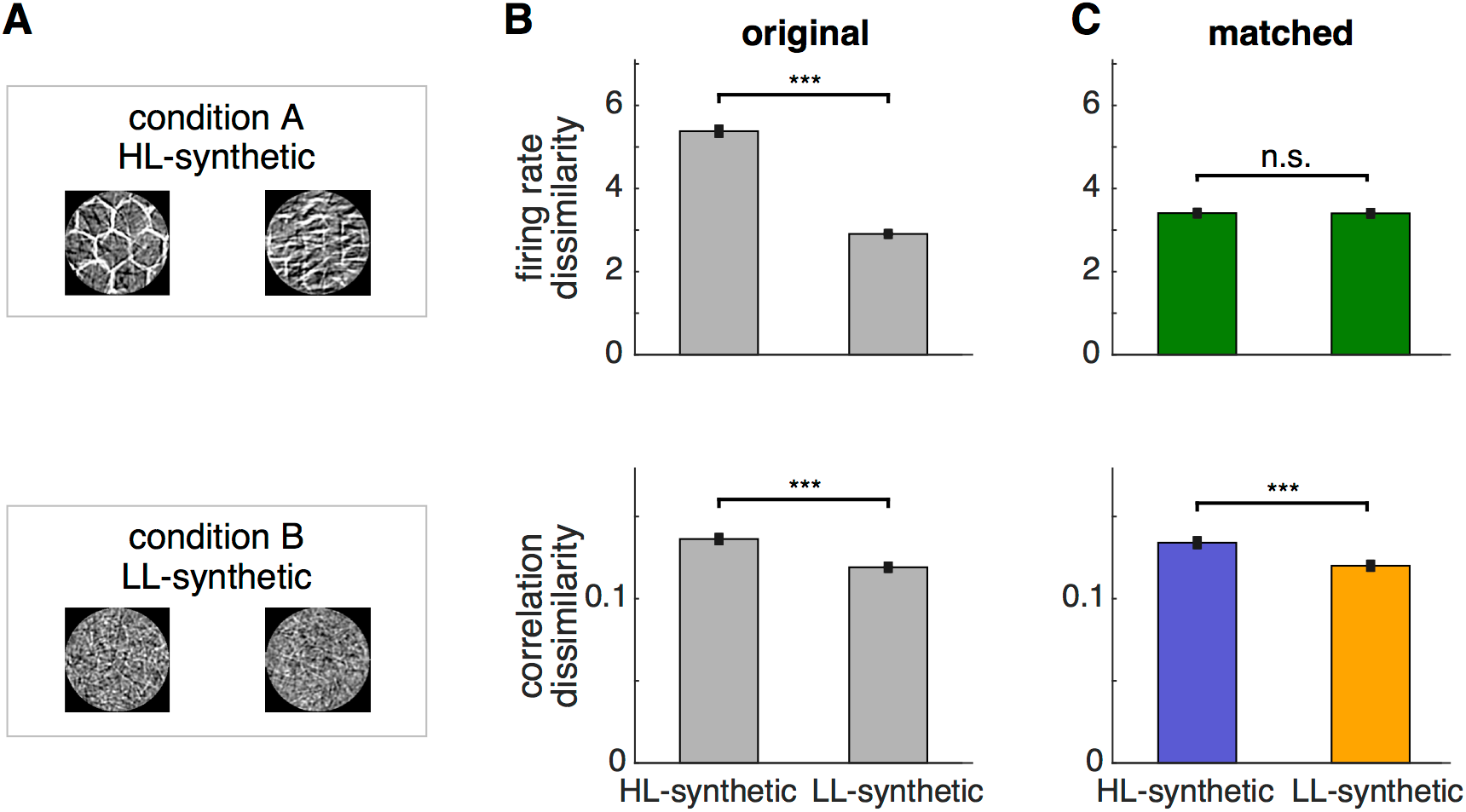
Comparison of stimulus-specificity of correlation patterns evoked by stimuli with different levels of statistical structure. **a**, condition A, a set of synthetic image patches in which filter co-occurrences define a texture structure (HL-synthetic stimuli), condition B, a set of synthetic image patches generated from a V1 model of images (LL-synthetic stimuli). **b**, Stimulus-specificity of firing rate responses (top panel) and spike count correlation patterns (bottom panel) in the original (unmatched) data. While correlations show higher specificity for HL-synthetic images, specificity of firing rate responses is also higher in the first condition. **c**, Firing and correlation dissimilarities after applying the contrastive rate matching procedure. Contrastive rate matching eliminates stimulus-specificity of firing rate responses, but the residual dissimilarity of spike count correlations is still significantly higher for HL-synthetic stimuli than that for LL-synthetic stimuli.

Taken together, these results demonstrate that SCCs are stimulus-specific but this stimulus-specificity hinges upon the higher-order structure of stimuli: removing high-level structure reduces stimulus-specificity, while reintroducing such a structure in controlled synthetic images restores the stimulus-specificity of SCCs. These differential effects of stimulus statistics cannot be accounted for by stimulus specific variations in spike counts.

## Discussion

We recorded population responses from area V1 of awake, task-engaged monkeys, and investigated how the correlation structure of V1 activity depends on stimulus content. Crucially, our analysis established that upon presentation of natural scenes the fine structure of correlations was specific to the presented stimulus. Furthermore, by designing synthetic image patches in which the statistical structure could be controlled, we demonstrated that the stimulus-specificity of SCCs was dependent on stimulus complexity: stimuli characterized by low-level structure showed reduced stimulus-specificity in SCCs, while images characterized by both low- and high-level structure showed increased stimulus-specificity. We argued that the stimulus-dependence of SCCs is a natural consequence of feedback in the ventral stream: inferences about high-level structure of images provide context for the interpretation of low-level structure through feedback influences, involving both lateral and top-down connections. We showed that a probabilistic hierarchical model of perceptual inference predicts the qualitative changes in stimulus-specificity of SCCs.

Parallel recordings from multiple neurons permit the investigation of higher order statistics of neuronal responses. Hence, the assessment of spike count correlations, commonly addressed as “noise correlations”, has become a central topic in neuroscience (Cohen and Kohn, 2011; Froudarakis et al., 2014; Kohn and M. A. Smith, 2005; Rikhye and Sur, 2015). Although measurement of spike-count correlations only requires the assessment of second-order statistics, accurate measurement (Cohen and Kohn, 2011; Ecker et al., 2010) and interpretation of variations in SCCs (Bányai et al., 2017) proved to be challenging. Factors, which affect the outcome of measurements include experimental design (number of repetitions, stimulus properties, e.g. static vs. moving), behavior, state dependent variables (eye movements, cognitive states), fluctuations in dynamical state and response bias (undersampling and firing rate) (Cohen and Kohn, 2011; de La Rocha et al., 2007). In our experiments we adopted a task design that aimed to control for a number of these factors and we also introduced additional controls in the analysis to eliminate those confounds that task design could not eliminate. To obtain a reliable estimate of pairwise correlations, we used a paradigm that permits a large number of repetitions under controlled conditions. This enabled us to limit sample variance in our measurements. Although anesthesia can permit a larger number of repetitions and/or a larger stimulus set, the associated paralysis causes stereotyped relations between stimulus and RF locations that can introduce artificial correlation structures. Using awake and task-engaged monkeys eliminates this confound and ensures that collective fluctuations, which introduce uncontrolled factors into the measured correlations, are minimized (Ecker et al., 2014). Eye movements have been shown to contribute to correlations in the visual cortex (McFarland et al., 2016). In our case monkeys were rewarded to fixate on a spot at the center of the screen during the presentation of off-foveal stimuli and trials in which fixation could not be maintained were removed from the analysis. However, during fixation, microsaccades and slow-drift eye movements could still occur and consequently, they could introduce collective changes in responses. While saccades represent voluntary motor actions and are known to be affected by stimulus-content (Meermeier et al., 2016), microsaccades and eye-drifts, are largely considered involuntary, have random (exponentially distributed) onset times and thus, are unlikely to affect the comparison of correlation similarity across conditions. Major other factors contributing to changes in spike count correlations in V1 have recently been identified on the basis of a large dataset (Schulz et al., 2015). These factors are cortical distance, tuning similarity, firing rate, spike isolation and spike width. Of these factors, only firing rate is changing across the conditions that we contrasted in our experiments. Therefore this potential confound required special attention. We developed the contrastive rate matching procedure and demonstrated its power on synthetic data before applying it to physiological data. This analysis confirmed that our conclusions on stimulus-dependence of correlations were not a consequence of changing firing rates.

Recently, stimulus-dependent modulations in pairwise correlations were observed in vitro in retinal ganglion cells (Franke et al., 2016; Zylberberg et al., 2016). These studies demonstrated that the microcircuitry of the retina can introduce stimulus-specific correlations that are precisely tuned to enhance decodability of sensory signals. However, these in vitro preparations are limited to the simple circuitry of the retina and leave open the question on hierarchical processing of complex stimuli in the cortex. To balance extra variability in neural activity in awake behaving animals, instead of restricting ourselves to pairs of neurons we analyzed a population of neurons. Here, characterization of the full correlation matrix was central to adequately assess the stimulus-specificity of SCCs. Earlier studies in mice used two-photon calcium imaging to characterize the changes in correlation structure as a result of changes in stimulus statistics (Hofer et al., 2011; Rikhye and Sur, 2015). These studies assessed the correlated variance in the activity of pairs of neurons, as reflected by the calcium signal, across repeated presentations of the same stimulus. Stimuli were presented in long windows and were either movies or a sequence of moving or static gratings, thus responses to individual stimuli were not analyzed. These results extended earlier observations that stimulus-statistics affects higher-order single-cell response statistics (Froudarakis et al., 2014; Haider et al., 2010). While these results suggested that the fine structure of SCCs might be specific to stimuli, calculating noise correlations for stimuli that were changing during any given trial prevented the assessment of stimulus-specificity of correlations.

Our approach can be regarded as an extension of earlier work investigating the patterns of mean responses in the hierarchy of the visual cortex (Freeman et al., 2013). It has been shown that variance in mean responses in V2 can be well predicted by variations in factors that determine the statistics necessary for the generation of natural textures, while variations in mean responses in V1 can be predicted by variations in statistics at the level of independent Gabor-like edge filters. Similarly, contextual modulation of V1 activity by top-down influences from V2 neurons was demonstrated when high-level inferences were made in artificial images (Klink et al., 2017; Lee and Nguyen, 2001). These results can be explained in terms of probabilistic inference in a hierarchical internal model of natural images, in which mean responses correspond to the most probable interpretation of the stimulus. Our results support a computational framework in which pairwise statistics of neuronal responses reveal that the uncertainty associated with the inferences is also represented.

Probabilistic computations require the representation of probability distributions, i.e. the representation of the whole array of possible interpretations of a stimulus rather than only the representation of the best interpretation. Representation of probability distributions was proposed to be achieved by stochastic sampling (Hoyer and Hyvarinen, 2003; Lee and Mumford, 2003), which interprets response variability as a direct consequence of perceptual uncertainty. The stochastic sampling framework generalizes naturally to hierarchical computations (Lee and Mumford, 2003). Recently, it was shown that stimulus-dependence of both membrane potential and spike count variability in V1 can be predicted by a model of natural images (Orbán et al., 2016). This model, however, lacked the hierarchical structure presented here and was therefore unable to account for stimulus-dependent changes in correlation patterns. However, it provided a simple, but important demonstration of contextual modulation of V1 responses: the assessment of stimulus-contrast for a complete image patch affected the interpretation of local image elements (Wainwright and Simoncelli, 2000). Such contextual modulation can be explained by an important computational component, divisive normalization (Carandini and Heeger, 2011; Schwartz and Simoncelli, 2001). Sampling has also been linked to task-related changes in spike count correlation patterns (Haefner et al., 2016; Lange and Haefner, 2016). Our results fit naturally in the sampling framework by assuming that neural activity patterns at any given time represent individual, multivariate samples from the probability distributions both at the level of V1 and higher-level areas, e.g. V2 (Fig. 1).

### Alternative interpretations

Patterns in higher-order statistics of neuronal responses beyond the mean activity, namely single-cell variability and SCCs, have been observed in association with task-related modulatory effects, such as those driven by attention (Ecker et al., 2016; Rabinowitz et al., 2015; Ruff and Cohen, 2014) and in association with stimulus-related modulatory effects, such as those occurring during the perception of natural scenes (Froudarakis et al., 2014; Haider et al., 2010; Rikhye and Sur, 2015). Computations underlying both of these processes invoke high-level inferences: inference of task variables in the case of attention and inference of high-level stimulus features, e.g. object identity, in the case of perception. In both cases, inference of high-level variables breaks the feed-forward processing hierarchy in the visual cortex and introduces top-down effects (Gilbert and Sigman, 2007). Such modulatory effects of top-down computations related to attention have been demonstrated in population responses throughout the visual processing hierarchy, both regarding single cell statistics (mean (Reynolds and Heeger, 2009) and variance (Goris et al., 2014)) and pairwise statistics (SCCs (Ecker et al., 2016; Haefner et al., 2016; Rabinowitz et al., 2015; Ruff and Cohen, 2014)). A recent phenomenological model of correlations suggested that multiplicative components might reflect top-down influences (Lin et al., 2015). Here, the so-called affine model could account for patterns in correlations emerging in anesthetized animals through collective gain modulation. However, stimulus-dependence and stimulus structure dependence cannot be addressed by this approach. Our result that correlation patterns depend on stimulus structure suggests that top-down influences involve processes more elaborate than simple collective gain modulations.

Alternative approaches have been proposed to understand the correlational structure of evoked and spontaneous activity using balanced networks (Hennequin et al., 2014b; Renart et al., 2010). Recently, it was proposed that patterns in correlations can be captured by a simple phenomenological model with feed-forward and lateral connections (Rosenbaum et al., 2016). Such approaches are complementary to the functional approach adopted in this study. In functional models of the visual cortex, parameters of the models are solely determined by the statistics of stimuli (Orbán et al., 2016). While there are attempts to link functional models to the network architecture (Aitchison and Lengyel, 2016; Hennequin et al., 2014a), the way neural circuits implement these computations is largely unexplored. Thus, studies on balanced networks provide useful constraints for determining plausible architectures.

Deep networks, which are characteristically hierarchical architectures for image processing, have become vastly successful in recent years, closing the gap between human and machine performance in complex visual tasks (LeCun et al., 2015). Prowess of deep learning architectures in tasks that humans excel inspired investigations into the parallels between biological visual systems and deep architectures (Kriegeskorte, 2015). These studies revealed structural similarities of the sensitivities of hierarchically organized neurons in the biological system and those in the deep learning model (Khaligh-Razavi and Kriegeskorte, 2014; Yamins and DiCarlo, 2016) and could also account for much of the variation in mean responses of inferotemporal units (Khaligh-Razavi and Kriegeskorte, 2014). The predominantly feed-forward architecture of these models shows impressive performance in a range of tasks but is still at odds with the biological system. Top-down influences of higher-level layers as well as recurrent connections within the same layer have no computational role in these models. In addition, these models do not feature an inherent representation of perceptual uncertainty. Thus, higher-order response statistics (including correlations) of neural responses are hard to reconcile with the working of classification-oriented deep networks. Congruent with the above observations, it has recently been shown that some of the discrepancies between performance of deep networks and humans in visual tasks seem to result from top-down interactions (Kar, K., Kubilius, J., Issa, E., Schmidt, K., and DiCarlo, J: Evidence that feedback is required for object identity inferences computed by the ventral stream. COSYNE 2017, Salt Lake City, Utah). These conclusions together with our results highlight the importance of understanding the function of feedback in shaping neuronal population responses and its role in behavior.

### Experimental procedures

#### Electrophysiological recordings

This study was conducted on two adult rhesus macaques (*Macaca mulatta*; Monkey A, male 8y and Monkey I, female, 12y). All experimental procedures were approved by the local authorities (Regierungspräsidium, Hessen, Darmstadt) and were in accordance with the animal welfare guidelines of the “European Union’s Directive 2010/63/EU”. We recorded extracellular signals (local field potentials (LFPs) and multi-unit activity (MUA)) from V1 using a chronically implanted microdrive containing 32 independently movable glass coated Tungsten electrodes with impedance between 0.7-1.5 MΩ and 1.5 mm inter-electrode distance (SC32, Gray Matter Research; Markovitz et al. 2011). The recording chamber was positioned based on stereotactic coordinates derived from MRI and CT scans following Paxinos et al., 2008. Signals were amplified (TDT, PZ2 preamplifier) digitized at a rate of 25 kHz and band-pass filtered between 0.1 and 300 Hz for LFP analysis and 300 – 4000 Hz for MUA recordings. For MUA analysis a threshold was set at 4 SD above noise level to extract spiking activity.

#### Behavioral paradigm

Animals were seated in a custom made primate chair at a distance of 64 cm in front of a 477x298 mm monitor (Samsung SyncMaster 2233RZ, 120 Hz refresh rate). Eye-tracking was performed using an infrared-camera eye-control system (ET-49, Thomas Recording). At the beginning of each recording week, the receptive field locations and orientation preferences of the recorded units were mapped with a moving light bar drifting in a randomized sequence in 8 different directions. The two monkeys performed an attention-modulated change detection task. To initiate the trials, the monkey had to maintain fixation on a white spot (0.1° visual angle) presented in the center of a black screen and press a lever. After 500 ms, two visual stimuli appeared in an aperture of 2.8-5.1° at a distance of 2.3-3.2° from the fixation point. One of the stimuli covered the receptive fields of the recorded units, the other was placed at the mirror symmetric site in the other hemifield. After an additional 700 ms, the color of the fixation spot changed, cuing the monkey to direct its covert attention to one of the two stimuli. When the cued image was rotated (20°), the monkey had to release a lever within a fixed time window (600 ms for monkey A, 900 ms for monkey I) in order to receive a reward. A break in fixation (fixation window 1.5° diameter) or an early lever release resulted in the abortion of the trial, which was announced by a tone signal. The number of completed trials varied between 524-1110 per recording session. No more than one session was recorded on a given day. In order to obtain a balance between the reliable estimation of spike count correlations (Fig. 3D) and the number of comparisons between stimulus pairs, we used 6 or 8 different stimuli per session, resulting in 65-180 repetitions (124 on average) per stimulus. The order of stimulus presentations was randomized. The number of good channels varied between 15 and 23 per session. Trials in which the signals were contaminated by clear electrical artifacts, were discarded from the analysis. The maximum number of trials discarded from a recording session was 3 (0.8 on average).

#### Visual stimulus design

Stimuli were either black and white natural images or synthetic images generated from an image model. Stimuli were presented in a square or circular aperture. We generated synthetic control stimuli that matched the low-level statistical properties of the natural images, but lacked any high-level statistical structure. As neurons in V1 are sensitive to oriented edges, we designed a set of 3000 Gabor filters adapted to the receptive field characteristics of the recorded neurons and these filters were linearly combined to obtain a synthetic image patch. For each control stimulus, we sampled the activations of 500-3000 Gabor filters from the empirical distribution of filter responses to the corresponding natural image. The pixel distributions of control images were then matched to the corresponding natural ones in terms of mean (luminance) and variance (contrast). For a second set of experiments, we reintroduced higher-level statistical structure to synthetic images by calculating the responses of the filter set on photos of natural texture patterns, then setting up a correlation matrix for filter activations in such a way that two filters were more strongly correlated if their responses to the texture photo were more similar. Sampling from this correlated filter activation distribution resulted in texture-like synthetic patterns corresponding to the statistical structures typically represented in V2. In each recording session, half of the stimuli were synthetic images with statistical structure corresponding to the representation in V1 (LL-synthetic stimuli), and the other half consisted of natural and synthetic images with structures corresponding to representations in V2 (HL-synthetic stimuli), in the first and second set of experiments, respectively.

#### Dissimilarity of population responses

In each recording session, evoked responses on each of the D electrodes and in each trial were characterized by a spike count calculated in a 400 ms window 360 ms after stimulus presentation. The delay was introduced in order to eliminate transients in neuronal responses that distort measurements of spike rate correlations due to stimulus locking. V1 population responses to a particular stimulus were characterized by (i) a firing rate vector of dimension D, obtained as the normalized mean spike count of each channel over the set of trials presenting the given stimulus, (ii) a spike count correlation matrix of dimension DxD, calculated between the z-scored spike counts over the same trial set. The dissimilarity between the population responses to two specific stimuli can thus be calculated in terms of firing rates and correlations. For firing rates, we used the L2 distance between the firing rate vectors. For correlations, we used the averaged absolute difference between specific pairwise correlation values. This kind of dissimilarity measure was selected over distance measures calculated between whole matrices, such as the KL divergence, in order to allow for the control of correlation dissimilarities caused by firing rate dissimilarities. The KL divergence quantifies the dissimilarity between two zero-mean, equal-variance Gaussian distributions, with covariance matrices equal to the measured correlation matrices. For all statistical comparisons and assertions of significance, unpaired t-tests were used.

#### Baseline for dissimilarity

Assessment of response statistics, including mean and correlations, requires repeated presentation of the same stimulus. The limited number of repetitions results in variance of the measures, including the measures for rate and correlation dissimilarities. A baseline for dissimilarity can be established by randomly splitting the trials for a given stimulus into two sets and comparing the measures on the two sets of samples. Evolution of self-dissimilarity can be assessed for different numbers of repetitions and the trend can be extrapolated to high trial numbers. We fitted rate and correlation dissimilarities at low trial numbers to predict those at high trial numbers and found that a double-exponential function fitted best the dependence of dissimilarity on trial number (data not shown). We used this extrapolation to establish a lower bound on the dissimilarity values in analyses where across-stimuli dissimilarities were established for both of the compared conditions.

#### Contrastive rate matching

Spike count correlations and firing rates are not independent due to the nonlinear mapping from membrane potential fluctuations to spiking activity (de La Rocha et al., 2007). Consequently, dissimilarities in correlations can also depend on dissimilarities in firing rates between responses evoked by different stimulus pairs. In order to establish the stimulus-dependence of correlations that is independent of changes in firing rates we constructed a measure that controls for firing rate changes. First we constructed a pairwise measure of firing rate. We chose the geometric mean rate, GMR, as it has been shown to predict correlations well (Schulz et al., 2015) (we implemented the control with the arithmetic mean firing rate as well and obtained similar results, data not shown). Instead of model-based controls, such as inferring membrane potential correlations (Dorn and Ringach, 2003) or just normalizing the correlations by GMRs (Kohn and M. A. Smith, 2005) (we implemented the latter but it did not alter the conclusions of our paper, data not shown), we implemented a non-parametric control to compare correlation dissimilarity under two conditions. Distributions of GMR differences were matched in the two conditions, similar to the matching of firing rate distributions when comparing response variability in Churchland et al (2010) (Churchland et al., 2010). To do so, we pooled all channel pairs from all sessions related to one condition (e.g. natural images), and calculated both GMR differences and correlation differences between all stimulus pairs for all channel pairs. The two conditions yield two joint distributions of GMR and correlation differences. To equate GMR differences in the two conditions we construct a marginal distribution of GMR differences using identical bins for the two distributions, take the minimum of the number of samples in matching bins, and subsample the data points in the condition with the higher sample count. Thus, we obtain the same GMR difference distributions in the two conditions and correlation differences can be compared on the GMR difference matched data sets (see Fig. 4A for a visual description of the process).

#### Controlling for finite data effects

As spike count correlations are calculated from a finite number of trials of finite length, increasing the number of spikes can also increase correlations without any additional top-down effects. Since our predictions concern exactly these top-down effects, we want to exclude the possibility that bottom-up differences together with finite measurements can account for the observed differences. We applied the raster marginal model (RMM) to control for this confound (Okun et al., 2012), which can be used to define a probability distribution over spike count correlation matrices, and is parametrized by spike counts on individual channels, and spike counts in individual time bins. For both natural images and V1-level synthetic images in each session, we sampled 500 correlation matrices from the RMMs defined by the evoked responses to each stimulus in the two conditions. From the simulated correlations we calculated the distributions of mean dissimilarities for the two conditions. Thus, we obtained an estimate of how much marginal spiking statistics pre-determine the differences in dissimilarities observed in the data, and how much of those is attributable to effects not accounted for by the RMM.

#### Validation of controls using simulated neural activity

Concomitant changes in firing rates and correlations make it challenging to isolate the effects of the stimulus on one or the other. We therefore devised a network of neurons where the two variables can be manipulated separately, which enabled us to validate that controls (e.g. contrastive rate matching) are efficient in separating the effects of rate and true correlation modulation. We simulated a population of 40 neurons, for which the mean and correlation of membrane potential activities were sampled from Gaussian and LKJ (Lewandowski et al., 2009) distributions, respectively. These parameterized a multivariate Gaussian distribution that was used to obtain membrane potential samples in 20 ms time windows (1 ms when a full raster is produced to specify a raster marginal model). Thus, variabilities of membrane potential samples are dependent across the population but independent across time bins. A firing rate nonlinearity established instantaneous firing rates for individual cells and spikes were obtained tracking the integer crossings of the integral of the fluctuating firing rates. This model has been shown to reproduce both single-cell statistics (Carandini, 2004; Dorn and Ringach, 2003) and pairwise statistics of V1 neurons (Bányai et al., 2017). Variability in spiking responses is dominated by membrane potential variance since the spiking model introduces minimal additional variability. This setting has two important characteristics. First, it avoids the use of a Poisson variability since the firing rate nonlinearity ensures a linear relationship between mean firing rate and variance but circumvents the washout of membrane potential covariability from spiking statistics by excess private variability (Bányai et al., 2017). Second, it expresses the characteristic relationship between mean firing rate and spike count correlation (de La Rocha et al., 2007). The network of neurons was used to assess the effects of firing rate and correlation modulations under conditions that approximated those in physiological experiments.

### Supplemental Information

Supplemental Information includes Supplemental Experimental Procedures and three figures.

### Author contributions

A.L., L.K. and W.S. designed the experiments, M.B., A.L. and G.O. conceived the study, L.K. and J. K-L. trained the animals and recorded the data with technical assistance from A.L., M.B. designed stimuli, performed the analysis and simulations, M.B. and G.O. analyzed the results, all the authors discussed the results and wrote the paper.

## Acknowledgements

We are grateful to Gareth Bland for providing data-management support. We thank Balázs Ujfalussy, Máté Lengyel, and József Fiser for comments on an earlier version of the manuscript. This work was supported by a Lendület Award (G.O.), the National Brain Program (NAP-B KTIA NAP 12-2-201, G.O.), the Deutsche Forschungsgemeinschaft (DFG NI 708/5-1, A.L.) and the European Union’s 7th Framework Programme (FP7/2007-2013 Neuroseeker, W.S.).

